# Characterisation and functional predictions of canine long non-coding RNAs

**DOI:** 10.1101/303966

**Authors:** Céline Le Béguec, Valentin Wucher, Lætitia Lagoutte, Edouard Cadieu, Nadine Botherel, Benoît Hédan, Clotilde De Brito, Guillory Anne-Sophie, Catherine André, Thomas Derrien, Christophe Hitte

**Affiliations:** Univ Rennes, CNRS, IGDR (Institut de génétique et développement de Rennes) - UMR 6290, F-35000 Rennes, France.; Centre for Genomic Regulation (CRG), The Barcelona Institute of Science and Technology, Dr. Aiguader 88, Barcelona 08003, Spain. Universitat Pompeu Fabra (UPF), Barcelona, Spain.; UMR PEGASE, Agrocampus Ouest, INRA, 35042, Rennes, France.

## Abstract

Long non-coding RNAs (lncRNAs) are a family of heterogeneous RNAs that play major roles in multiple biological processes. We recently identified an extended repertoire of more than 10,000 lncRNAs of the domestic dog however, predicting their biological functionality remains challenging. In this study, we have characterised the expression profiles of 10,444 canine lncRNAs in 26 distinct tissue types, representing various anatomical systems. We showed that lncRNA expressions are mainly clustered by tissue type and we highlighted that 44% of canine lncRNAs are expressed in a tissue-specific manner. We further demonstrated that tissue-specificity correlates with specific families of canine transposable elements. In addition, we identified more than 900 conserved dog-human lncRNAs for which we show their overall reproducible expression patterns between dog and humans through comparative transcriptomics. Finally, co-expression analyses of lncRNA and neighbouring protein-coding genes identified more than 3,400 canine lncRNAs, suggesting that functional roles of these lncRNAs act as regulatory elements. Altogether, this genomic and transcriptomic integrative study of lncRNAs constitutes a major resource to investigate genotype to phenotype relationships and biomedical research in the dog species.

## Introduction

With the advancement of high-throughput sequencing technologies, transcriptome analyses (RNA-seq) have made it possible to identify major RNA classes, including the long non-coding RNA class (lncRNA)^1,2^. The transcriptome thus corresponds to sets of transcribed RNA molecules, with or without the ability to code for proteins, for a given time, condition or tissue^3^. Arbitrarily defined according to a size criterion (generally greater than 200 nucleotides), lncRNAs possess similar characteristics than RNAs encoding proteins (mRNAs), i.e. they can be spliced and have (or not) a polyadenylation tail, but they are differentiated by a lack of a functional open reading frame. Following the sequencing of all the RNA transcripts, the annotation and classification of the different RNAs consist of reconstructed transcript models, from which it is crucial to define their functional roles. LncRNAs can be located either in intergenic regions (*lincRNAs*), often close to protein-coding genes, or overlapping the opposite strand of mRNAs (*antisense*) and, therefore, both represent strong candidate sequences that modulate transcription of nearby protein-coding genes.

The domestic dog has emerged as a relevant model for studying the genetic basis of numerous traits, including Mendelian and complex diseases, morphology, physiology and behaviour^4^. Genomic resources have expanded our understanding of the canine genome but exhaustive annotation of functional elements, including long non-coding RNAs, remains necessary to facilitate the identification of genotype-phenotype relationships^5^. Most recently, the number and types of canine lncRNAs have increased considerably^6,7^ however, little is known about their function and biological roles. To our knowledge, a few canine lncRNAs have been characterised experimentally, as illustrated by the lincRNA close to the *BAIAP2* gene linked to podocyte migration^6^ and the lncRNA *GDNF-AS* involved in a Hereditary Sensory Autonomic Neuropathy (HSAN) in hunting dogs^8^. As in other species, it is expected that lncRNAs in the dog genome are expressed at a lower level and display higher tissue-specificity than protein-coding genes. Also, because lncRNAs are subjected to rapid sequence turnover during evolutionary processes, they are seldom conserved in vertebrates^9^. In addition, lncRNAs are also known to regulate the expression of protein-coding genes through *cis*-acting or *trans*-acting regulation mechanisms. LncRNAs located near to and overlapping protein-coding genes have attracted particular attention from researchers and several studies in humans and mice have uncovered several mechanisms, including *cis*-regulation of the expression of their protein-coding neighbour and overlapping partners^10^.

Knowledge of the biological functions of lncRNAs is continuously increasing, but their origin and evolution are still poorly understood. One recent hypothesis that has emerged concerns the theory that transposable element (TE) sequences are a possible source of non-coding exons^11^. In humans and mice, TEs are frequently found in lncRNAs^12,13^ and numerous studies emphasise the importance of TEs in the regulation of gene expression (for a recent review see Chuong et al.^14^). In dogs, TEs also occupy a substantial fraction (40%) of the genome and might be important actors in the origin of functional novelties. More particularly, a well-studied family of TE is the specific TEs (SINEC_Cf)^15^, that have been implicated in many dog genetic diseases^16,17^ or phenotypic differences between dog breeds^18,19^.

In this study, we performed an exhaustive analysis of 10,444 canine lncRNA genes for which we describe the overall transcriptional profiles across 26 distinct tissue types, representing various anatomical systems. Analysis of their expression patterns retrieved relevant relationship between tissue types and showed evidence of tissue-specificity for a large fraction of lncRNAs. Co-expression analysis of lncRNAs and nearby protein-coding genes was performed to infer putative functionality of uncharacterised lncRNAs using the principle of ‘guilt by association’^20^ from their co-expression with genes of known functions. Together, we provide a large and unique resource that characterises the transcriptional patterns of lncRNAs, improves the knowledge of the origins of canine lncRNAs and contributes to infer their potential functionality.

## Results

### Landscape of canine IncRNA transcription

We produced 16 RNA-seq in a previous study^7^ and collected 10 published data^6^, corresponding to a total of 26 stranded RNA-seq data to represent a panel of diverse canine anatomical systems (Supplementary Table S1). We applied a state-of-the-art RNA-seq bioinformatics pipeline^21^ to assess gene expression levels for both canine lncRNAs and mRNAs (see the Methods section). For lncRNAs, we used the FEELnc-based classification^7^ to differentiate long intergenic non-coding RNAs (lincRNAs, n = 5,651) from lncRNAs that have overlapping protein-coding genes in antisense (antisense, n = 4,793). We determined that a total of 7,763 (74.3%) lncRNAs are expressed in at least one tissue with a normalised count^22^ TPM > 1 where TPM = Transcripts Per Million and 9,542 (91.4%) with a TPM > 0.1. By comparing their respective expression levels with protein-coding genes mRNAs, canine lncRNAs have, on average, 20 times lower expression levels (Wilcoxon test, p-value < 2.2^e-16^) for all tissues, with the notable exception of the testis tissue for which the difference is 6 times less pronounced, a trend also showed in other species^2,23^ (Fig. 1a). We then compared the expression of lncRNAs in various tissue types and found that the mean expression level in testis tissue is much higher (TPM = 7.02) than in any other tissue (ranging from 0.11 for the pancreas tissue to 1.60 for the hair follicle tissue and with an average TPM = 1.05) (Fig. 1a). By comparing lincRNA versus antisense expression patterns (Supplementary Fig. S1), we observed that lincRNAs have a lower expression than antisense, except for testis tissue (Wilcoxon test, p-value < 2.2^e-16^).

**Figure 1.**
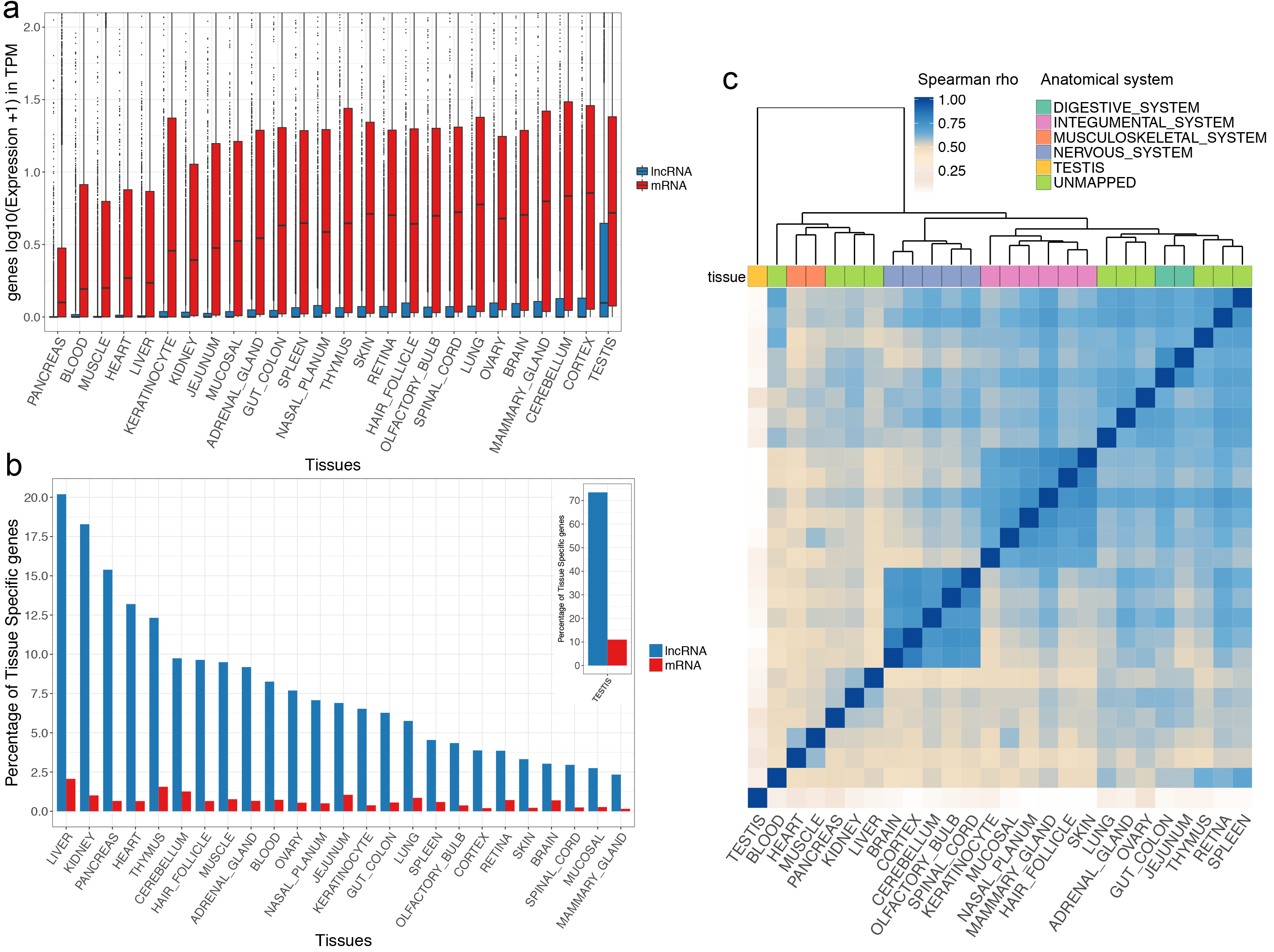
(a) Comparative analysis of log10 transcription (TPM + 1) levels between mRNAs (red) and lncRNAs (blue) genes in 26 canine tissues. (b) Proportions of tissue-specific lncRNAs (blue) and mRNAs (red) amongst expressed genes in each tissue with testis tissue represented in the separated panel (top right) (c) Hierarchical clustering of 26 tissues based on Spearman correlations measured from lncRNA expression data.

Taking advantage of the variety of samples, we analysed whether canine lncRNAs displayed patterns of tissue-specific expressions by calculating a tissue-specific score ‘tau’^24^. This score, ranging from 0 to 1 (with tau = 1 being a highly tissue-specific expression), has been shown to be highly robust when evaluating tissue-specific genes^25^. Using a stringent threshold ≥ 0.95, a total of 44% of lncRNAs (n = 4,599, Supplementary Table S2) displayed a clear pattern of tissue-specificity, highlighting the potential specialised functionality and distinct spatial pattern of expression in the corresponding tissue. In comparison, only 17% of mRNAs (n = 3,635) showed a pattern of tissue-specificity reflecting that, as in other species, canine lncRNAs are more tissue-specific than mRNAs^26^. Among all classes of lncRNAs, we observed that lincRNAs are more tissue-specific (68.6%) when compared to antisense (31.4%) (Chi-square test, p-value < 2.2^e-16^). As shown in humans, the canine testis tissue is particularly enriched in tissue-specific lncRNAs^1,2^ (n = 3,001). This highlights the singularity of this tissue, probably due to the presence of many cell types, the state of its chromatin and the binding of specific transcription factors^27^. Excluding testis tissue, we identified an average of 63 lncRNAs expressed specifically by tissue. In order to assess the relative enrichment of tissue-specific lncRNAs versus tissue-specific mRNAs, we measured the proportion of tissue-specific genes amongst the total number of expressed genes (Fig. 1b). As an example, this highlighted that 20% of lncRNAs expressed in liver tissue are liver-specific, compared to only 2% of mRNAs. As previously shown, testis tissue is unusual, with 73% of the lncRNAs expressed being testis-specific as against 11% for mRNAs.

Given that tissue-specificity can be influenced by cells with common origins, we then investigated the tissue-specificity within tissues related to the nervous system including brain, cerebellum, cortex, olfactory bulb and spinal cord. We found specific expression patterns for 260 lncRNAs in the different samples of the nervous system^28^. Among the nervous system samples, we identified 156 lncRNAs specifically expressed in the cerebellum, 60 in the cortex and 44 in the olfactory-bulb (Supplementary Table S2); (tau threshold ≥ 0.95). These lncRNAs were characterised by a higher expression level (mean TPM = 3.30) in comparison to other tissues (mean TPM = 0.12). For example, we found a cerebellum-specific lincRNA (tau = 0.96), annotated in humans as *CASC18*, with a role in neural cell differentiation^29^.

Although we identified a large number of tissue-specific lncRNAs, we then clustered the 26 tissue samples based on their lncRNA expression profiles in order to detect groups of lncRNAs that exhibit common expression patterns. The heatmap in Fig. 1c highlights that clustering the lncRNAs’ expression data recovers biologically meaningful relationships between tissue types. This analysis defined two main clusters with more than 3 samples, which grouped nervous system tissues (n = 5) and integumental system tissues (n = 6) and supported the fact that some lncRNAs shared expression patterns in multiple tissues. This analysis was repeated using only mRNA expressions and showed similar clusters (Supplementary Fig. S2). Complementary to the previous tissue-specific analysis, this clustering approach allowed us to identify an lncRNA (*RLOC 00033166*) expressed in all nervous system tissues with a mean TPM = 7.89 but not detected in any other tissues. Interestingly, this lncRNA is transcribed in antisense orientation to the Neuregulin 3 gene (*NRG3*), which is involved in neuroblast cell differentiation^30^ and thus represents a potential candidate for its regulation. Another lncRNA (*RLOC_00020746*) is almost only expressed in the musculoskeletal system cluster (heart and muscle) and is expressed in antisense to the Popeye Domain Containing 3 gene (*POPDC3*), which may play an important role in the development of skeletal muscle and heart tissues^31^. Apart from the clustered groups of tissues identified in the analysis, most samples, and particularly the testis sample, remained strongly unrelated to any other tissues.

### Tissue-specific expression of lncRNAs correlates with their transposable element content

TEs are thought to constitute part of the active lncRNA regions^11^ and may drive the specific expression patterns observed for lncRNAs^14^. Interestingly, in dogs, a canine-specific SINE repeat family (SINEC_Cf) has been shown to provide many polymorphic site insertions^15^ and is also associated with multiple disease phenotypes^16,18^. To gain insights into the relationships between canine lncRNAs and transposable elements, we analysed the TE content of the 10,444 lncRNAs in the 4 major TE classes (DNA transposons, LTRs, LINEs and SINEs) as annotated by the RepeatMasker database^32^. We first showed that almost 70% of canine lncRNA transcripts contain at least one TE overlapping an lncRNA exon. We then determined that ∼20% of the cumulative exon sequences of lncRNAs are composed exclusively of TEs (Fig. 2a).

**Figure 2.**
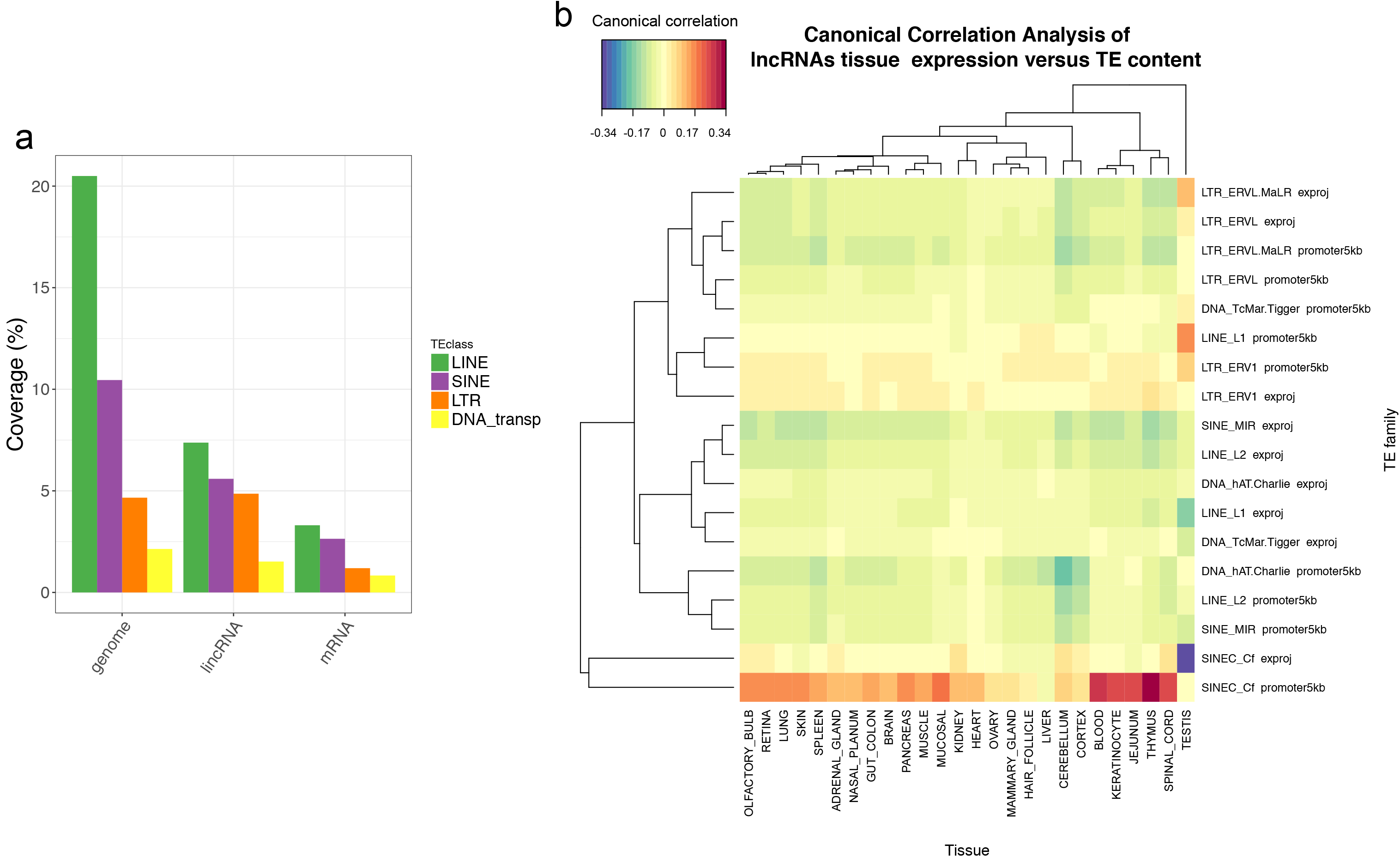
(a) Proportion of transposable elements (LINEs, SINEs, LTRs and DNA transposon in green, purple, orange and yellow respectively) with respect to the entire dog genome, lncRNA exons and mRNA exons (from left to right). (b) Canonical correlations between lncRNAs’ expression in the 26 tissues (column) and TE families in exons and promoters (row).

This proportion of lncRNA exonic TEs is less than that observed in the entire dog genome (37.7%) but is 2.5 times higher than for mRNAs exonic sequences (7.9%), highlighting the high prevalence of TEs in dog lncRNA exons as also observed in other species^13^. Compared to the genome-wide distribution of TEs, most TE families are under-represented in lncRNAs with the exception of LTR-ERVL retroviruses (Supplementary Fig. S3), which are also significantly enriched in human lncRNAs^13^. In addition, we computed canonical correlations between TE content in both lncRNAs exons and promoters and lncRNA tissue expressions using the mixOmics program^33^ (see Methods). This analysis highlighted that lncRNA expression in the different tissues tends to be associated with high contents of SINEC_Cf specifically in lncRNA promoters (Fig. 2b), a pattern which is not seen for mRNAs (Supplementary Fig. S4). These results suggest that changes in the promoter sequences of lncRNAs by inserting TEs and, more particularly SINEC_Cf, may contribute to the specific spatial expression of canine lncRNAs. The analysis also pinpoints that exons of lncRNAs expressed in testis are particularly depleted in SINEC_Cf, suggesting that testis-specific expression of canine lncRNAs is more possibly associated with other family of TE such as LTR (Fig. 2b) and probably LTR-ERVL (Supplementary Fig. S3) as recently highlighted in mouse germline^34^.

### Identification of dog-human conserved lncRNAs by comparative genomics

DNA conservation among divergent species is a widely used indicator to infer functionality and to suggest conserved functions.Identifying orthologous lncRNA relationships is challenging since lncRNAs are more likely to be gained or lost during evolution^9^ than constrained sequences as protein-coding genes. Therefore, we used a positional comparative genomics approach, parameterised at the synteny level, to predict putative lncRNA orthologues between dog and human using the EnsEMBL Compara database^35^ (see Methods). Using this strategy, we mapped the positions of canine lncRNAs onto the human genome and identified 939 lncRNAs (9.0%) with 1:1 human non-coding orthologues from the GENCODE database^36^ which can be separated into 560 lincRNAs (9.9% of the total number of lincRNAs in dogs) and 379 antisense (7.9% of the total number of antisense in dogs) (Fig. 3 and Supplementary Fig. S5). We found that lncRNA orthologues that were part of a synteny block shared similar gene structures, with a mean number of exons per gene of 3.6 and 3.1 for humans and dogs, respectively (Supplementary Table S3). Based on the GENCODE annotation of the conserved dog-human lncRNAs, we were able to confirm the annotation of the well-studied *HOTAIRM*^37^, *MALAT*^38^, *NEAT_1*^39^ and *PCA3*^40^ in dogs and to infer novel canine orthologous relationships such as the CASC family of lncRNAs (*CASC15, CASC17, CASC18* and *CASC20*) and *INHBA-AS1*^41^ or *MEG9*^42^. Complementary to this synteny-based strategy, we evaluated the selective constraints acting on canine lncRNAs and mRNAs by using the Genomic Evolutionary Rate Profiling (GERP) scores^43^ (see Methods). In comparison to mRNAs, we showed that the 939 syntenically conserved lncRNAs are significantly less constrained than syntenic mRNAs (Wilcoxon test, p-value = 9.54^e-171^). In addition, this allowed us to pinpoint that syntenically conserved lncRNAs tend to be more constrained (mean GERP = 0.221) than non-syntenic lncRNAs (median GERP = 0.195), although not statistically significant (Wilcoxon test, one-tailed p-value = 0.19). Altogether, this is in agreement with other studies that have reported a lower purifying selection acting on lncRNAs^9^.

**Figure 3.**
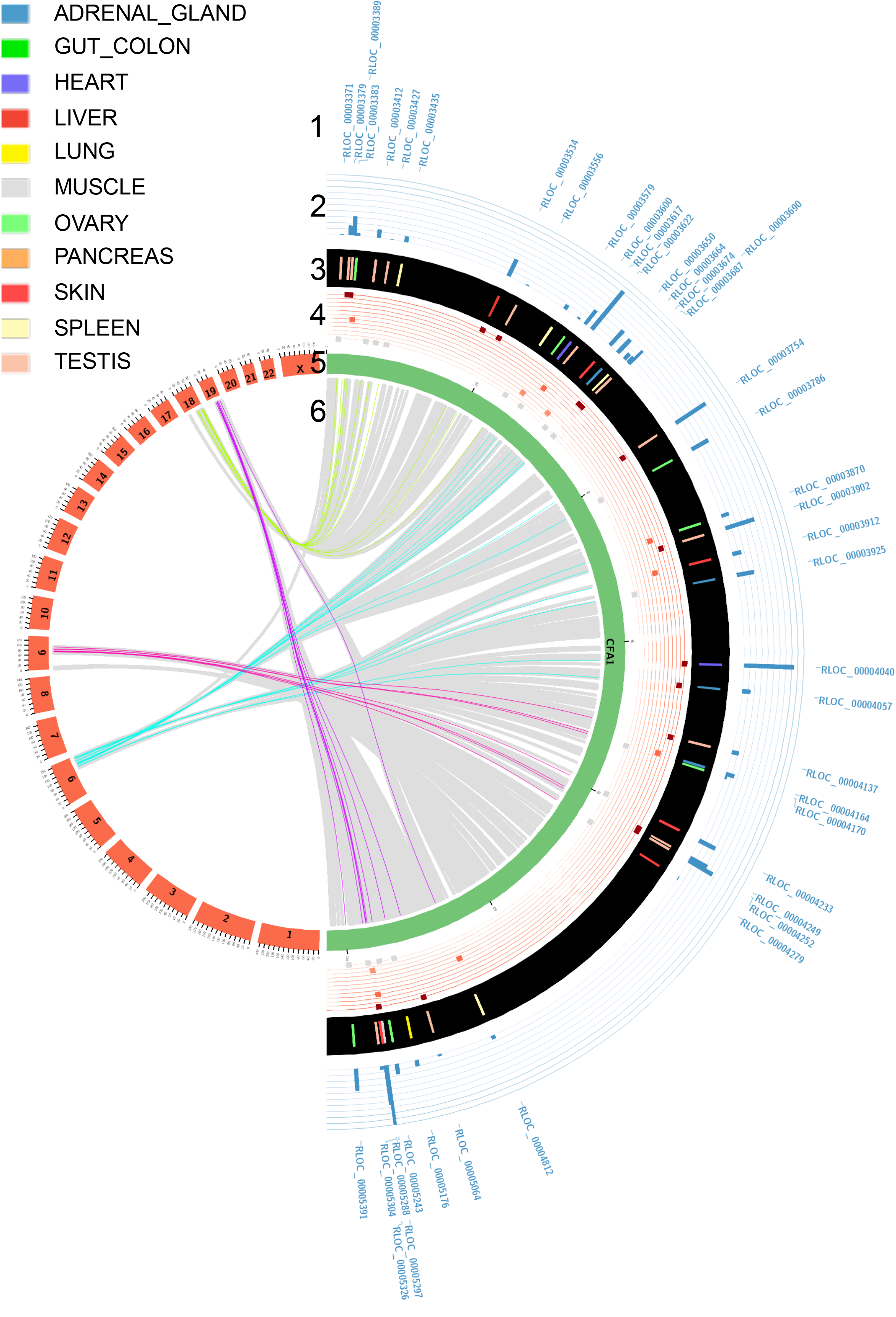
The Circos plot provides the visualisation of dog-human orthologous relationships of lncRNAs and their comparative transcriptomic-based expression patterns. For canine chromosome 1, tracks are described from the outside to the inside. Track1: Labels of canine lncRNAs identified with human orthologues are described at the most outside track of the figure. Track2: Level of expression of canine lncRNAs is shown by the blue histogram. Track3: The tissue in which lncRNA is expressed in dogs. The 11 tissues are adrenal gland, gut-colon, heart, liver, muscle, ovary, pancreas, skin, spleen and testis and are represented with the colour code as shown in the legend. Track4: Expression of the human orthologous lncRNA is represented. 11 lines are reported. When the human orthologue lncRNA is expressed in the same tissue and with the highest of expression, it is represented by a dark red square on the upper line. When an orthologous human lncRNA is expressed in the same tissue, with the second maximum of abundance, it is represented on the second line in lighter red. When an orthologous human lncRNA is not expressed in the same tissue, it is represented by a grey square. Track5: The green layout depicts the canine chromosome, the red layout represents the human chromosomes. Track6: For the innermost part, coloured lines link the dog-human orthologous relationships of lncRNAs. Grey lines represent the orthologous relationships of protein-coding genes.

### Analysis of transcriptional profiles of conserved lncRNAs through comparative transcriptomics

As sequence conservation is not an infallible indicator of functionality, we integrated the expression patterns of orthologous genes in our analysis. It has now been reported that orthologous genes may have conserved or variable expression profiles between species such as in human and mouse^44^. To investigate the expression patterns of orthologous lncRNAs between dogs and humans, we determined the correlation of the level of expression between 11 human tissues from the ENCODE project^45^ that matched the dog tissues. We produced bar charts (Fig. 4) to represent the expression levels of each conserved lncRNA in the 11 different samples of both species and these allowed us to visualise and compare expression patterns of orthologous lncRNAs between dogs and humans (Supplementary Fig. S6). Overall, of the 727 conserved lncRNAs with expression in at least one tissue on both species, mean lncRNA expression patterns correlate well between dog and human, such that expressed genes in dogs tend to be expressed in the same tissues in humans (mean correlation rho = 0.39). However, among these general tendencies, we could identified a subset of 26% of dog-human orthologous lncRNAs (n = 192) with significantly similar expression patterns (p-value < 0.05; mean rho = 0.87), suggesting that these lncRNAs might be involved in evolutionary conserved functions. In addition, these lncRNAs are mostly tissue-specific (average tau = 0.94 and 0.93 for dogs and humans, respectively), a finding already reported between human and mouse where tissue-specific lncRNAs in one species are also tissue-specific in other^44^. These evolutionary and transcriptionally conserved lncRNAs are mostly expressed in testis (n = 154) then spleen (13), lung (9), liver (8), skin (5), and heart (4) (Fig. 4a). Anatomical systems such as the brain, testis, heart, liver and kidney were previously found to have clearly distinct signatures of tissue-specific genes in humans and mice^44^ and showed strong conservation between the two species. Here, as an example, the lncRNA *TRDN-AS1*, transcribed in the opposite strand of the Triadin gene (*TRDN*), is found specifically expressed in the heart tissue in both human and dog species. Another example is given by the *L1NC01698* gene which exhibits a shared expression pattern between dog and human in the skin, as described using GTEx data portal^46^. We also defined lncRNAs (n = 77) with intermediate correlation pattern (p-value < 0.2; mean rho = 0.50), which display a more tissue-variable expression between species (Fig. 4b). This subset shows that sequence constraints do not imply an exact conserved expression and that different factors in the two species may act on the expression of the same gene, modulating its expression level. A third subset of lncRNAs (n = 458) was defined with divergent transcriptional pattern (p-value > 0.2; mean rho = 0.17) between the two species (Fig. 4c). This subset may indicates expressed lncRNAs in anatomical systems with complex structures composed of many primary cell types, for which tissue functionality may differ between samples and/or between species because of their substructure.

**Figure 4.**
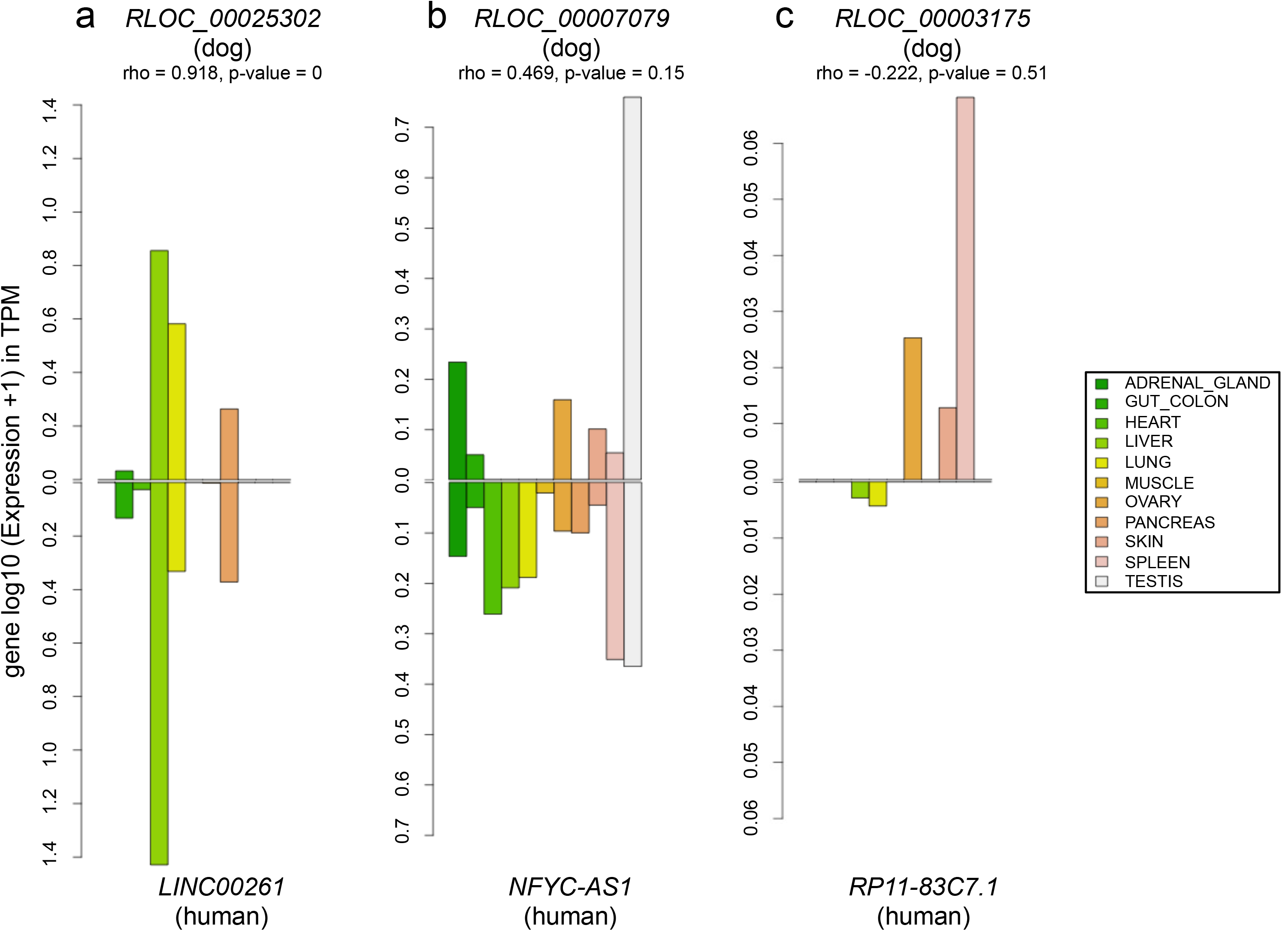
Bar charts representing the expression of the 3 lncRNAs in 11 matched dog-human tissues, (a) high conservation of expression, (b) intermediate conservation of expression, (c) low conservation of expression.

### LncRNA:mRNA co-expression and correlation analysis

In many species, lncRNAs have been involved in regulating gene expression by coordinating epigenetic, transcriptional or post-transcriptional processes^47^. Co-expression pattern analyses provide a means to investigate sequences that modulate the transcription of nearby genes. In addition, regulatory sequences are often located near to their target gene, modulating their transcription. Hence, identifying correlated expressed sets of protein-coding and non-coding loci provides a way to (i) identify relationships between lncRNAs and mRNAs and (ii) infer potential regulatory functionality of lncRNAs.

Here, we searched for significant co-expression, computed through statistical correlations between lncRNAs and their neighbouring protein-coding genes. We used the ‘guilt-by-association’^20^ strategy to functionally annotate the lncRNA co-expressed with its mRNA neighbour, especially in the case of divergent transcripts sharing bidirectional promoters^48^. We retained neighbouring protein-coding genes transcribed in divergent and convergent orientations (excluding same strand) with respect to the lncRNA in order not to bias the correlation analysis with lncRNA being actually unannotated UTRs of the neighbour protein-coding genes. On average, these canine lincRNAs are located at 25 kb of the closest protein-coding genes. We identified more than 126,000 lncRNA:mRNA pairs within a 1 Mb window using the FEELnc classifier module^7^. We then divided pairs into two classes consisting of antisense:mRNA pairs, defined by lncRNAs overlapping mRNAs transcribed in antisense (n = 3,401) and lincRNA:mRNA pairs defined by intergenic lncRNAs located at less than 1 Mb of an mRNA (n = 123,456). Using the expression data from the 26 tissues, we computed expression correlations and mined the resulting pairs. We found 8,139 significant correlations (| rho | > 0.5 and p.adjust BH^49^ < 0.05) consisting of 7,615 lincRNA:mRNA and 524 antisense:mRNA pairs for a total of 3,410 distinct lncRNAs. These results revealed co-expressed pairs that may predict regulatory relationships. Among significant correlations, only 58 pairs were found with a significant negative correlation coefficient (rho < −0.5). The lncRNA co-expressed in these pairs are significantly more distant to their mRNAs (mean distance = 471 kb) compared to the ones in the positive correlations (mean distance = 394 kb) (Wilcoxon test, one-tailed p-value = 0.02), indicating that they may be located in different genomic regions. An interesting example of positive correlation is given by the lincRNA *RLOC_00018074* co-expressed with the mRNA *EGFR* (Epidermal Growth Factor Reception) in the 26 tissues (Fig. 5a). *RLOC_00018074* and *EGFR* genes are transcribed in a divergent orientation and, by using published data which mapped dog promoters from H3K4me3 marks^50^ (Fig. 5b), we could hypothesise that this pair share a bi-directional promoter thus reinforcing the validity of our functional inference.

**Figure 5.**
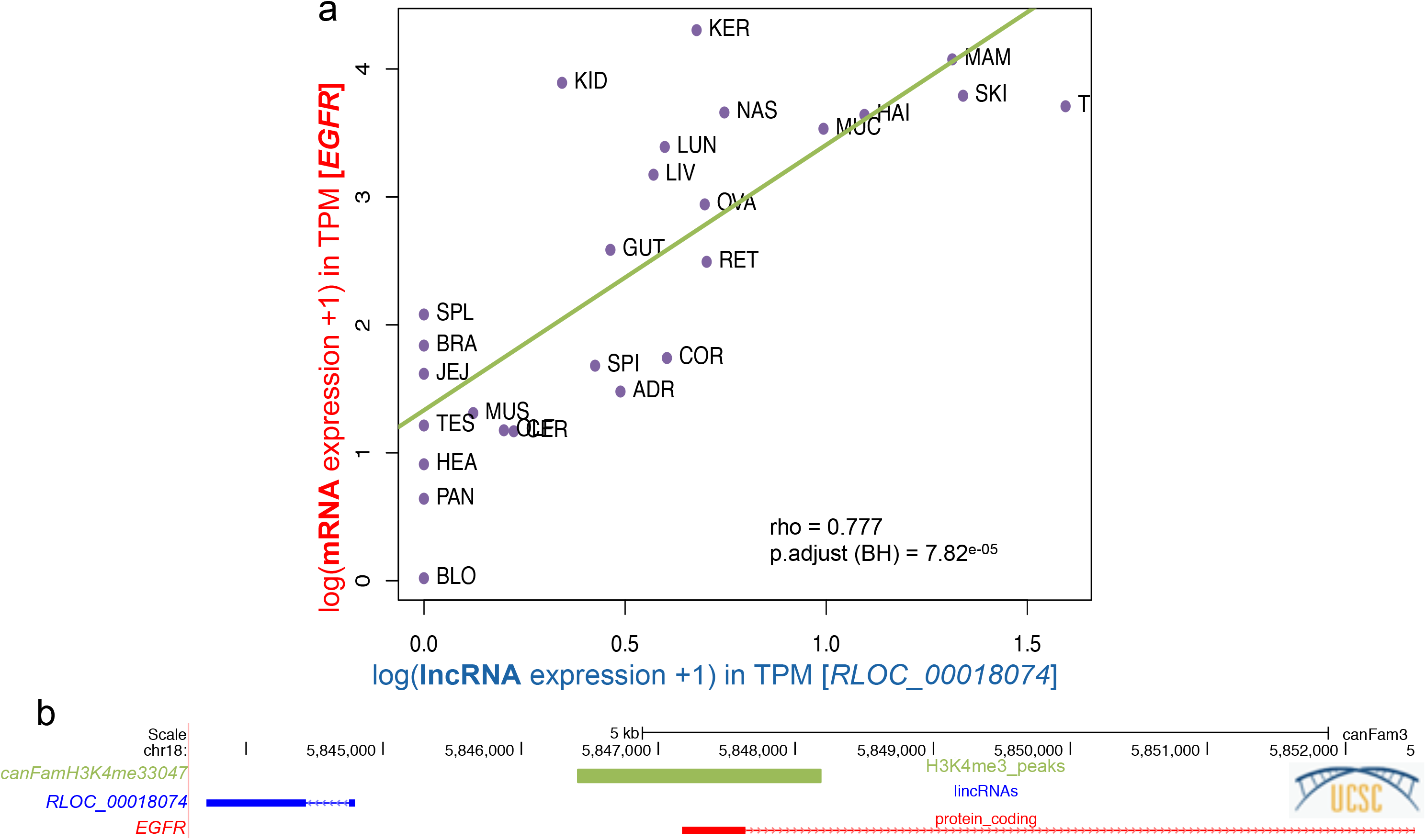
(a) Correlation plot of the lncRNA:mR\Apair (*RLOC_000018074:EGFR*) in the 26 tissues, (b) UCSC screenshot representing the genomic localisation of the *RLOC_000018074:EGFR* pair transcribed in a divergent orientation with the promoter mark (H3K4me3) from published data.

Of the 3,410 lncRNAs, we observed multiples type of co-expression patterns. A subset of 1,691 (49.6%) lncRNAs have two or more mRNAs significantly co-expressed for which more than 800 (55%) displayed a higher correlation with the second nearest protein-coding gene or greater (Supplementary Fig. S8). It shows the importance of identifying co-expression patterns for establishing accurate lncRNA:mRNA pairs that may interact. In addition, the strength of the correlation decreases with the distance between the two elements of the pair. Indeed, highly co-expressed gene pairs (1 > | rho | > 0.9) are closer (average = 89 kb) than less correlated co-expressed gene pairs (0.7 > | rho | > 0.5; average = 411 kb).

Finally, we performed a gene ontology (GO) term enrichment analysis in order to predict potential lncRNA functions. Based on the human orthologous mRNAs (n = 3,977) with which the lncRNAs are co-expressed, we focused our GO analysis on the Biological Process (see Method). A total of 22 GO terms were significantly enriched (adjusted p-value < 0.05) and correspond to developmental processes such as ‘sensory organ development’ (GO: 0007423, 147 genes), ‘axon development’ (GO: 0061564, 129 genes) or ‘hindbrain development’ (GO: 0030902, 49 genes) (Supplementary Table S4). These 22 GO terms included 913 unique mRNAs co-expressed with their corresponding canine lncRNA (26.8%). These results predict a functional assignment for these 913 canine lncRNAs which represent interesting candidates for studies related to the enriched biological processes.

## Discussion

Transcriptomic studies have emphasised the analysis of long non-coding transcripts using expression profiles to characterise the patterns and their potential functional roles. LncRNAs tend to be expressed at lower levels than protein-coding genes, being under less stringent evolutionary constraints, and are preferentially enriched in regulatory functions. Here, we realised a global analysis of long non-coding RNA expression across multiple tissues to aid genome annotation and improve functional annotation of the dog genome. By using strand-specific RNA sequencing of 26 tissues, we profiled the expression patterns of lncRNAs and achieved a detectable expression for 91% of lncRNA in at least one tissue, which constitutes the largest resource of long non-coding RNA expression data sets in dogs. We showed the higher tissue-specificity pattern of lncRNAs relative to mRNAs, and this suggests specialised functions in the development, differentiation and physiological processes of tissues. The largest number of tissue-specific lncRNAs was shown to be in the testis, a result that can be related to the pervasive transcription during spermatogenesis process and due to chromatin remodelling^51^.

There is growing evidence showing a close association of transposable elements (TEs) with non-coding RNAs. Thousands of long intergenic non-coding RNAs are associated with endogenous retrovirus LTR TEs in human cells^11^. Exhaustive characterisations of the links between lncRNAs and TEs are becoming fundamental as diseases and phenotypic traits are increasingly found to have a TE and/or lncRNA etiology^11,14^. Here, we have shown a strong association of the SINEC_Cf TE family with lncRNAs in dogs. The SINEC_Cf family have the capacity to spread in the genome^15^ and, therefore, may be able to introduce regulatory sequences into lincRNAs or their promoter sequences upon insertion. These results provide the first evidence for discovering how and which TE families represent a major force in shaping the canine lncRNA expression and their tissue-specificity expression patterns in a lineage-specific fashion.

Despite the number of tissues investigated in this study (n = 26), tissue-specificity is a limiting factor in annotating the whole repertoire of lncRNAs of a species, since their annotation depends on the availability of numerous tissue and cell types as well as multiple spatio-temporal conditions. Additional analysis of other tissues and development stages will be needed for identifying an extended catalogue of lncRNAs and their isoforms. Until now, most expression studies have been performed on a population level usually averaging the transcriptomes of millions of cells^52^. Pioneering RNA-seq of single cells has provided the characterisation of transcriptional differences of both coding and non-coding RNAs on a genome-wide scale^53^. The single-cell RNA-seq approach, a relatively recent experimental approach, will be able to detect and characterise the expression patterns of complex tissues, in order to provide high-resolution identification of cell types and markers and to identify splicing patterns and allelic random expressions that are variable between cells. Single-cell RNA-seq will help in-depth, functional characterisation and gene expression variation derived from different genetic backgrounds at the cell resolution.

Co-expression analyses rely on similarities in gene expression and are widely used for the functional annotation of unknown genes using the principle of ‘guilt-by-association’^20^. Focusing on neighbouring genes, we tested the expression correlation of lncRNA:protein-coding pairs and observed that thousands of lncRNA-coding pairs showed an overall positive correlation, suggesting coordinated transcription and, by implication, a shared function or pathway. Whilst constructing co-expression networks is rather straightforward, the resulting network of connected lncRNA:mRNA do not provide information on the nature of the regulatory relationship of the connected genes, which limits its biological interpretation. In our co-expression analysis, weak interactions or interactions only occurring within a single, or very few, tissue types might have been missed through Spearman correlation and an under-estimate of the numbers of coordinated transcriptions and potential *cis*-regulations in which lncRNAs might be involved.

Next-generation sequencing technologies have considerably advanced the field of comparative genomics and comparative transcriptomics. These approaches are particularly important for studying the evolution of gene regulation in model organisms, investigating the level of the lncRNA sequence and expression conservation with humans and investigating the way that this conservation determines conditions in which the dog constitutes an appropriate model for diseases and phenotypic traits. Using a comparative genomic approach, we identified more than 900 lncRNAs with a human orthologue and the analysis of the level and distribution of expression between dog and human allowed us to define conserved or diverged expression patterns that then served to predict putative, conserved functions or to pinpoint more subtle changes of functionality. Indeed, tissues are complex structures that are composed of many primary cell types and it is unclear whether tissue functionality remains similar. It is not yet known to what extent the relative expression of genes has been conserved among the species or whether the tissue samples in the different species remain similar or differ in their substructure.

Much comparative data still needs to be produced and analysed concerning transcriptional changes associated with differentiation and development or with cellular responses to external stimuli. Indeed, the dog is a promising model for complex phenotypes, genetic diseases and clinical studies. Altogether, this genomic and transcriptomic integrative study of lncRNAs constitutes a major resource for the dog species.

## Materials-Methods

### Datasets

The 26 RNA-seq sample dataset (Supplementary Table S1) represents a wide variety of canine cell-types and tissue-types. The 26 tissues can be classified by Anatomical System as inspired by the Expression Atlas database (https://www.ebi.ac.uk/gxa/home/), as shown in the Supplementary Table S1. Two tissues correspond to ‘Digestive system’, 6 tissues to ‘Integumental system’, 2 tissues to ‘Musculoskeletal system’ and 5 tissues to ‘Nervous system’. The others tissues correspond to ‘Unmapped’ and ‘Testis’.

RNA extraction and directional sequencing were performed as described in Wucher et al.^7^ and Hoeppner et al.^6^ and are available through accession numbers SRP077559, SRX111061 - SRX111071, and SRX146606 - SRX146608.

The canine ‘canFam3.1-plus’ annotation^7^ (containing 10,444 lncRNA and 21,810 protein-coding genes (mRNAs)) was used as the reference annotation for this study and the canFam3 assembly version was the reference genome.

### Quantification of mRNAs and lncRNAs expressions

Based on the bio-informatic protocol described in Djebali et al.^21^, FASTQ reads were aligned on the canine reference genome and transcriptome using the STAR^54^ program (version 2.5.0a) and we determined the gene and isoform expression levels for both lncRNAs and mRNAs for each of the 26 RNA-seq with RSEM^22^ (version1.2.25). Finally, the RSEM output files were parsed in order to extract gene expression level, normalised in Transcripts Per Million (TPM) and merged to obtain one matrix expression file with gene names in rows and tissue expression levels (TPMs) in columns.

### Characterisation of lncRNA expression

#### Tissue-specificity

To calculate the tissue-specificity score ‘tau’^24^, ranging from 0 to 1 (with tau = 1 for highly tissue-specific genes), we used the matrix file described above and calculated the following equation^24^:

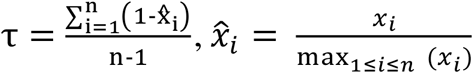

where *n* corresponds to the number of tissues analysed and *x_i_* is the gene expression in the tissue *i*. The specificity score was calculated using a threshold of 1 TPM (to be stringent).

To extract a stringent set of tissue-specific genes, we used a tau threshold greater than 0. 95 corresponding to a ratio of 4 between the first and the second tissue (e.g. max(*xi*) / max^2nd^(*xi*) > 4) or, in others words, corresponding to cases where gene *i* is four times more expressed in the first tissue compared to the second most highly expressed tissue. Note that we provide a perl script dedicated to computing the ‘tau’ score on GitHub available here: https://github.com/tderrien/IGDR/blob/master/script/specificityscore.pl.

#### Heatmap and clustering analysis

For each biotype (lncRNAs and mRNAs), we computed all pair-wise Spearman correlations between the 26 tissues in order to obtain a matrix of distance. Using this file, together with a meta-data file containing anatomical system classification (e.g. nervous system, integumental system, musculoskeletal system, digestive system, testis, and unmapped), the heatmap and the associated hierarchical ascendant clustering (with default parameters: euclidean distance and method = ‘complete’) were created as in Breschi et al.^44^ using R software (version 3.2).

#### TE content

Canine TE annotations were based on the RepeatMasker^32^ database, downloaded from the UCSC table browser^55^ (September 2017). Four major classes were analysed: DNA transposons and retrotransposons, LTRs, LINEs and SINEs; each of them being divided into families (e.g. SINEC_Cf). To compute the proportion of TEs per gene, exons of each gene were projected onto the genome and the number of exon-projected nucleotides overlapping TE families was computed using the bedtools program (version 2.19.0) on bed12 input files with the `-wao` and `—split` options. For gene promoters, the same process was repeated using 5 kb upstream of each gene starts.

#### TE content and gene expression

The mixOmics tool^33^ (version 6.3.1) was used to compute canonical correlations to highlight potential correlations between two matrices *X* and *Y*, where *X* is the matrix of lncRNA (or mRNA) expressions in the 26 tissues and *Y* is the matrix of TE content in lncRNAs (or mRNAs) exons and promoters. The `rcc` function of the mixOmics package (with method =’ridge’ and ncomp = 3) was used to seek for linear combinations of the variables (canonical variables) to reduce the dimension of the datasets. The heatmap was constructed with the mixOmics function `cim` (Clustered Image Maps) with default values, except ncomp = 3.

### Comparative genomics and transcriptomics

#### Comparative genomics

To identify conserved canine lncRNAs and mRNAs in humans, we used the Ensembl Compara database^35^. Briefly, the program annotates orthologous regions between two or more species based on Whole Genome Alignments (WGA) computed using the EPO (Enredo-Pecan-Ortheus) pipeline^35^. All canine gene coordinates were then mapped onto the human genome (version: GRCh37 using version 75, GENCODE version 19), having the same biotype (coding or non-coding) and considering only the one-to-one (1:1) relationships.

### GERP score

We used the EnsEMBL Compara API to retrieve GERP scores^43^ of the dog genome assembly (canFam3) (one wig file per each dog chromosome) computed on the 53 eutharian mammals. To get a single value per lncRNA and mRNA gene, we projected all exons of each gene onto the dog genome and calculated the median GERP score for each gene.

#### Comparative transcriptomics with human ENCODE data

We extracted the human gene expression using data produced by the ENCODE project^45^ (https://www.encodeproject.org/matrix/?type=Experiment&replicates.library.biosample.donor.organism.scientific_name=Homo+sapiens&biosample_type=tissue&assay_title=total+RNA-seq&assembly=hg19&award.project=ENCODE).We only selected data where tissues between humans and dogs could be matched. We found 11 dog:human matching tissues: adrenal gland, gut-colon, heart, liver, lung, muscle, ovary, pancreas, skin, spleen and testis. From the available RSEM files, we extracted the expression levels at the gene transcript levels in TPM for the dog data. When multiple RSEM files for a tissue replicates, we averaged the expression level per gene. We then determined the statistical correlation, with the Spearman correlation test, of the level and the distribution of the expression between these 11 human and dog tissues. Bar charts were created with R software to help visualise the comparison.

#### Comparative genomics and transcriptomic visualisation>

We used the Circos tool (circus-0.69-6) to visualise relationships of orthologous genes between dog and human. Circular layouts were constructed to allow global but meaningful figures with a Circos ideogram per dog chromosome.

### Co-expression analysis

#### Co-expression

To study the correlations of expression between lncRNAs and mRNAs, we used the FEELnc classifier module^7^ to annotate lncRNA classes and sub-classes (e.g. genic versus intergenic and divergent versus convergent, Supplementary Fig. S9) and to automatically identify all lncRNA:mRNA pairs in a sliding window of 1 Mb around each lncRNA. For the sake of interpretability, we filtered subclasses so that they only retained exonic antisense, intergenic divergent and intergenic convergent. For all lncRNA:mRNA pairs, we computed Spearman correlations between expression vectors (26 points) and corrected, associated p-values for multiple testing using the Benjamini-Hochberg (BH) method^49^. To identify significant pairs in co-expression, we used a correlation coefficient of | rho | > 0.5 and p.adjust < 0.05.

#### GO term analysis

From the 5,711 canine mRNAs that were significantly co-expressed with their lncRNAs (n = 3,410), we extracted human, orthologous EnsEMBL gene IDs of mRNAs (n = 3,977). To identify gene ontology (GO) terms significantly enriched in this list of gene IDs, we used the web-based GEne SeT AnaLysis Toolkit (WebGestalt^56^) software with the following settings: *Method of Interest* = ‘Overrepresentation Enrichment Analysis’ (ORA) and *Functional Database* class = geneontology (GO) named ‘Biological_Process_noRedundant’. Finally, we selected the *Gene ID type* ‘ensemble_gene_id’, to upload our orthologous data using ENSG IDs and selected genome for *Reference Set for Enrichment Analysis*. We can add ‘Advanced parameters’, so we keep default parameters (*Minimum Number of Genes for a Category*: 5; *Maximum Number of Genes for a Category*: 2000 and *Multiple Test Adjustment*: BH) and use the *Significant Level*: FDR 0.05.

All data are available through the dedicated website: http://dogs.genouest.org/lncRNA.html.

## Acknowledgments

The biological samples were obtained from the ‘Cani-DNA_CRB’, which is part of the CRB-Anim infrastructure (http://dog-genetics.genouest.org). We would like to thank the following funding agencies: the ‘Cani-DNA_CRB’ [ANR-11-INBS-0003], the CNRS (Centre National de la Recherche Scientifique), the Université Rennes1 and the Brittany Region (France) (PhD funding for CLB). We thank the LUPA consortium (http://eurolupa.org) and the BROAD Institute for the sequencing. We thank Sarah Djebali of the INRA GenPhyse of Toulouse and members of the canine genetics team at IGDR-Rennes. We thank Ignacio Gonzalez for help with the analysis using mixOmics.We also thank the Genouest Bioinformatic platform, Rennes, France (https://www.genouest.org/) for providing the necessary computing resources.

## Author contributions

C.L.B., T.D. and C.H. conceived the study and wrote the manuscript. C.L.B., V.W., L.L., E.C., B.H., N.B., C.D.B., A.S.G., C.A., T.D., and C.H. provided the samples and tissues for phenotyping. Data analysis was performed by C.L.B., V.W., T.D. and C.H. All authors read, improved and approved the manuscript.

